# Honey bee trypanosomatid parasite dynamically changes the transcriptome during the infection of honey bee and modifies the host physiology

**DOI:** 10.1101/529321

**Authors:** Qiushi Liu, Jing Lei, Alistair C. Darby, Tatsuhiko Kadowaki

## Abstract

Although there are many honey bee pathogens/parasites, it is still not understood how they change their gene expression to adapt to the host environment or how the host simultaneously responds to pathogen/parasite infection by modifying its own gene expression. Such interactions must lead to changes in the physiological states of both host and parasite. To address this question, we studied a trypanosomatid, *Lotmaria passim*, which can be cultured in medium and inhabit the honey bee hindgut. We found that *L. passim* dynamically modifies the expression of mRNAs associated with protein translation and the electron transport chain to adapt to the anaerobic and nutritionally poor honey bee hindgut at early stages of infection, and to become dormant at late stages of infection. Meanwhile, several genes are continuously up- or down-regulated during infection, including *GP63* as well as genes coding for host cell signaling pathway modulators (up-regulated), and those involved in detoxification of radical oxygen species as well as flagellar formation (down-regulated). *L. passim* infection only slightly increases honey bee mortality and does not affect the number of microorganisms in the gut microbiota; but it induces honey bee innate immune response. Upon infection, the host appears to be in poor nutritional status, indicated by the increase in the levels of mRNAs for *take-out* and *facilitated trehalose transporter* and the decrease of *vitellogenin* mRNA level. Simultaneous gene expression profiling of *L. passim* and honey bee during infection provided insight into how both parasite and host modify their gene expressions. This study presents one of the best models to understand host-parasite interactions at the molecular and cellular levels in honey bee.

## Introduction

Honey bees (*Apis mellifera*) play a significant role in agricultural crop production and ecosystem maintenance. Nevertheless, the number of managed honey bee colonies has dramatically declined across North America and Europe since 2006. Although there are many potential causes for the observed declines, pathogens/parasites are considered major threats to the health of honey bees (Evans and Schwarz 2011a, Goulson et al 2015). There are different kinds of honey bee pathogens/parasites, such as viruses, bacteria, fungi, protozoans, and mites (Evans and Schwarz 2011b). The honey bee host response to infections has been previously characterized (Doublet et al 2017). For instance, apoptosis seems to be an important response to microsporidian infections and the Toll and Imd immune signaling pathways respond to viral infections. However, the consequences of eliciting and maintaining immune responses on honey bee physiology have not yet been fully understood. Furthermore, we poorly understand how parasites adapt to the honey bee environment when they start establishing the infection and how they react against the host responses during infection maintenance. Elucidating these processes will provide critical insight into understanding honey bee (host)-parasite interactions.

Two *trypanosomatidae* species, *Lotmaria passim* and *Crithidia mellificae*, were shown to infect honey bee. *C. mellificae* was first identified in Australia in 1967 (Langridge and McGhee 1967). Recently, a novel trypanosomatid parasite was discovered and named *L. passim* (Schwarz et al. 2015). *L. passim* seems to be more prevalent than *C. mellificae* (Arismendi et al. 2016, Cavigli et al. 2016, Cepero et al. 2014, Cersini et al. 2015, Morimoto et al. 2013, Ravoet et al. 2014, Ravoet et al. 2015, Regan et al. 2018, Schmid-Hempel and Tognazzo 2010, Schwarz et al. 2015, Stevanovic et al. 2016, Vavilova et al. 2017). Although the association of *L. passim* infection with winter mortality of honey bee colonies was suggested in several studies (Ravoet et al. 2013, Runckel et al. 2011), the effects of *L. passim* infection on honey bee health and colony survival remain poorly understood. *L. passim* can be cultured in medium and specifically infects the hindgut when orally introduced to the honey bee (Schwarz et al. 2015). These characteristics are similar to those of *Crithidia bombi*, a major trypanosomatid parasite of bumble bee (Lipa and Triggiani 1988, Schlüns et al. 2010). *C. bombi* infection dramatically reduces colony-funding success, male production and colony size (Brown et al. 2003). Moreover, it has been shown that *C. bombi* infection impairs the ability of bumble bees to utilize floral information (Gegear et al. 2006). Bumble bee was shown to up-regulate several genes in immune signaling pathways, such as *Spatzle, MyD88, Drosal, Defensin, Hymenoptaecin* and *Apidaecin* in the early stages of *C. bombi* infection (Riddell et al. 2011, Riddell et al. 2014, Schlüns et al. 2010). However, changes in *C. bombi* gene expression profile during infection have never been studied. Since both *L. passim* and *C. bombi* are specifically present in hindguts of honey bee and bumble bee, respectively, they must interact with the gut microbiota. In fact, there is a negative correlation between the infection level of *C. bombi* and the relative abundance of *Apibacter, Lactobacillus* Firm-5 and *Gilliamella* bacteria in the gut microbiota of bumble bees (Cariveau et al. 2014; Mockler et al 2018). Yet, a recent study has shown that pretreatment with *Snodgrassella alvi* makes normal honey bees more susceptible to *L. passim* infection under natural conditions (Schwarz et al. 2016). A relatively simple honey bee gut microbiota shapes the gut microenvironment not only lowering the oxygen but also the pH level. Furthermore, the microbiota utilizes the indigestible compounds of pollen to produce short-chain fatty acids and organic acids which help to maintain the nutritional status of the honey bee (Zheng et al. 2017). *L. passim* may affect the functions of the honey bee gut microbiota.

The genome of *L. passim* was sequenced and contains approximately 9,000 protein-coding genes which are transcribed as polycistronic mRNAs, like in other trypanosomatid species (Runckel et al. 2014). This demonstrates that the expression of mRNAs is primarily controlled by post-transcriptional mechanisms such as *trans*-splicing and polyadenylation (Clayton 2002). It is thus possible to profile the gene expression of *L. passim* during its infection in honey bee by transcriptome analysis. In this study, we characterized honey bee-parasite interactions using *L. passim* and *A. mellifera* as a model system. Cultured *L. passim* cells were orally introduced to newly emerged honey bees and the gene expression profiles of *L. passim* (parasite) and the honey bee (host) were simultaneously analyzed at early, middle, and late stages of infection. We will discuss how honey bees respond to *L. passim* infection, how their physiological states are modified, and how *L. passim* changes its gene expression to first adapt to the host environment and then to maintain the infection.

## Results

### Infection of the honey bee by *L. passim* and its effect on bee mortality

We infected newly emerged honey bees with 10^5^ *L. passim* by ingestion and maintained them with 50 % (w/v) sucrose under laboratory conditions. As shown in Fig. 1A, the number of parasites remained constant up to eight days and there was a dramatic increase at 15 and 22 days after infection. However, there was a large variation in the number of parasites among the infected individual honey bees. These results demonstrate that *L. passim* starts to actively proliferate in the honey bee hindgut between 8 and 15 days after infection.

**Figure 1.**
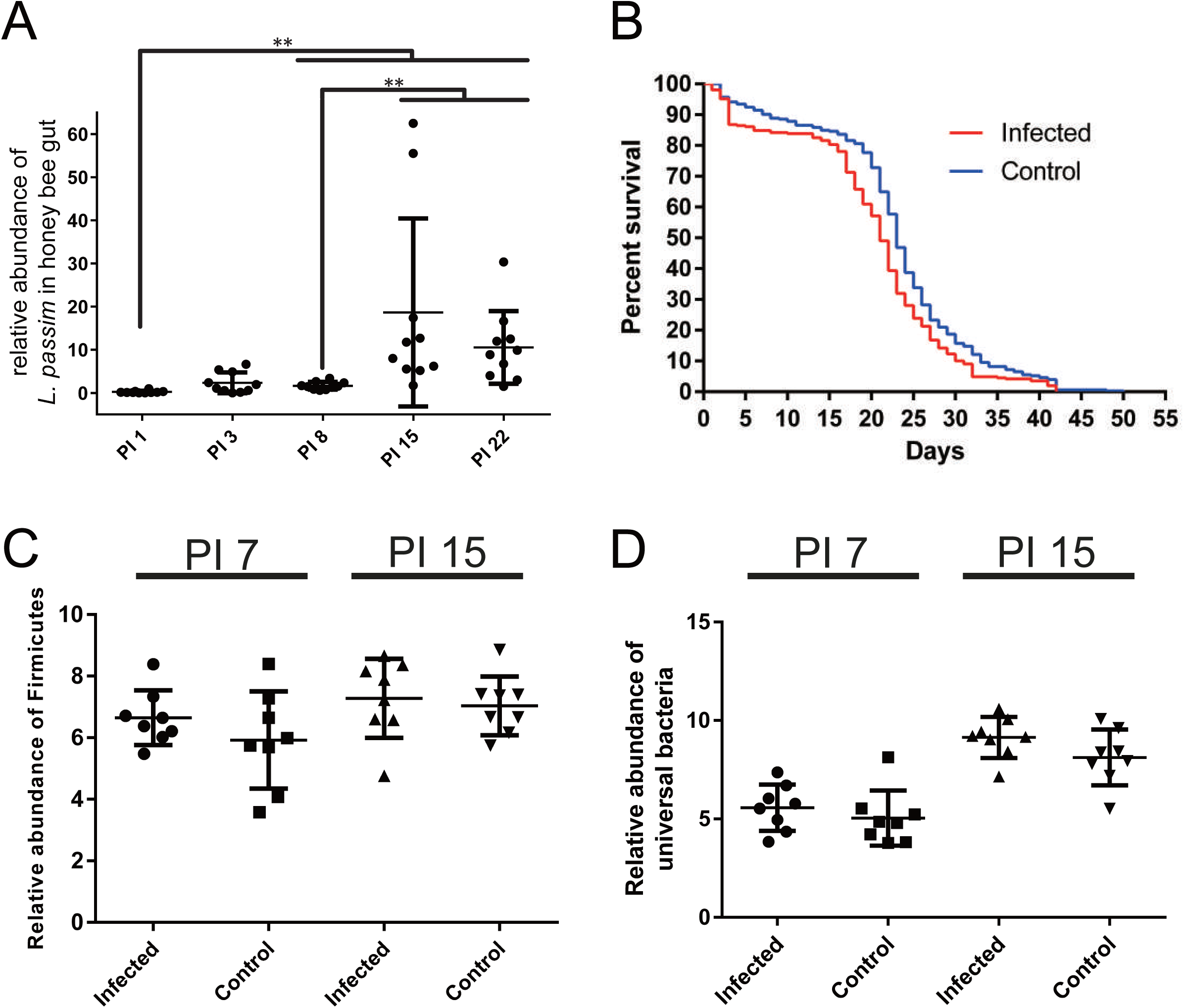
The effects of *L. passim* infection on honey bee mortality and gut microbiota. (A) The abundance of *L. passim* in ten individual infected bees at 1, 3, 8, 15 and 22 days after infection (PI 1-22). In order to measure the relative abundance of *L. passim*, one honey bee sample on day 1 was set as 1 and nine more samples were statistically analyzed on the same day. Mean values ± SD (error bars) are shown. The Steel-Dwass method was used for statistical comparison between the different time points. ** *P* < 0.01. (B) Mortality of honey bees infected by *L. passim* at 33 °C under laboratory conditions. 100 newly emerged honey bees were either infected with 10^5^ *L. passim* (Infected, red) or fed sucrose solution (Control, blue). The experiment was repeated three times and the averages are shown. The data were analyzed by the Log-rank (Mantel-Cox) test (*P*-value = 0.0002). (C) Relative abundance of Firmicutes in *L. passim*-infected (Infected) and the uninfected control (Control) honey bees at PI 7 and 15. (D) Relative abundance of universal bacteria in *L. passim*-infected (Infected) and the uninfected control (Control) honey bees at PI 7 and 15. Mean values ± SD (error bars) are shown. Unpaired *t*-test (two-tailed) was used for statistical analysis.

We also tested the mortality of honey bees infected by *L. passim* (at 33 °C) and fed with 50 % (w/v) sucrose under laboratory conditions. The infected honey bees survived 26-42 days after infection and this time period was slightly shorter than the survival time of control uninfected honey bees (Fig. 1B). These results indicate that the accumulation of parasites in the honey bee hindgut does not induce a rapid death of the host under laboratory conditions.

### Effects of *L. passim* infection on honey bee gut microbiota

Because *L. passim* and the majority of the gut microbiota co-exist in the honey bee hindgut, they are likely to interact. To test for potential interactions, we infected two-day-old honey bees with *L. passim* and fed them with a mixture of sucrose and pollen under laboratory conditions. We then compared the abundance of universal bacteria and Firmicutes (*Lactobacillus* Firm-4 and Firm-5) in the whole guts of both parasite-infected and uninfected control honey bees at 7 and 15 days after infection. There were no significant differences between the two groups at any of the examined time points (Fig. 1C-D). These results suggest that *L. passim* does not dramatically affect the general landscape of the honey bee gut microbiota.

### Changes in the gene expression profile of *L. passim* in the honey bee hindgut

We infected newly emerged honey bees with *L. passim*, returned them to their original hive, and then collected the parasite-infected honey bees at 7, 12, 20, and 27 days after infection (post infection (PI) 7, 12, 20, and 27). The RNA expression profiles of *L. passim* were analyzed together with those of cultured parasites, by RNA-seq. The mapping rates of clean reads to the *L. passim* genome were 69-78 % except for PI 7 (16 %). These were consistent with the above results which demonstrate that parasite population increases 8 days after infection under laboratory conditions. Clustering analysis revealed that the gene expression profiles of *L. passim* change during infection in the honey bee hindgut, except for the PI 20 and PI 27 samples (Fig. 2A). Interestingly, those changes were most dramatic at PI 12 and it was an outgroup to the other time points (Fig. 2B). Pair-wise comparisons between cultured parasites and honey bee-infecting parasites indicated that 411, 961, 415, 582 genes were up-regulated and 451, 1069, 658, 832 genes were down-regulated at PI 7, 12, 20, and 27, respectively (Fig. 2C and D) (Supplementary Table 1). As expected from clustering analysis, genes specifically up- and down-regulated at PI 12 represent the highest fraction of total genes changed during parasite infection (1126, 34.0 %). The genes continuously up- and down-regulated from the middle to the late stage of infection (PI 12-27 and PI 20-27) were the second highest fraction (744, 22.5 %). The third highest fraction corresponded to the genes specifically up- and down-regulated at PI 7 (476, 14.4 %). 44 genes were continuously up-regulated and 67 genes were continuously down-regulated throughout the parasite infection (Fig. 2C and D). The list of continuously up-regulated genes was enriched with the following gene ontology (GO) terms: “modulation by symbiont of host protein kinase-mediated signal transduction,” “modulation by symbiont of host nitric oxide-mediated signal transduction,” and “proteolysis.” It included homologs of glycoprotein (GP) 63 (leishmanolysin) and many peptidases (Supplementary Table 2). Oxidoreductase, tryparedoxin 1, tryparedoxin-like, tryparedoxin peroxidase, and paraflagellar rod protein 5 are included in the 67 down-regulated genes (Supplementary Table 3). These results suggest that *L. passim* affects several honey bee signal transduction pathways to establish and maintain hindgut infection. Furthermore, the parasites also need to adapt to the anaerobic environment of the honey bee hindgut (Zheng et al. 2017).

**Figure 2.**
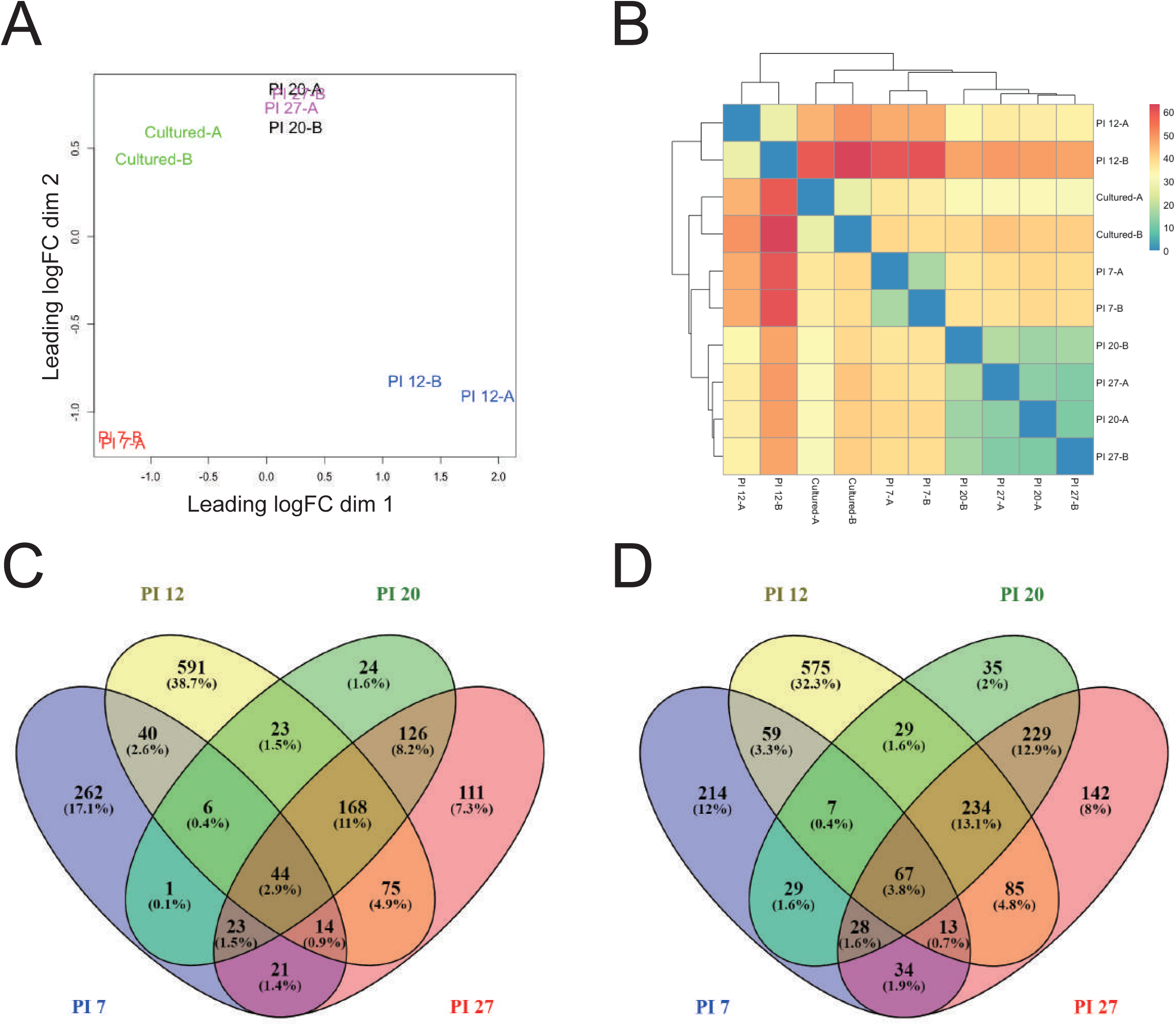
Global transcriptomic profiles of *L. passim* at different time points during infection in the honey bee hindgut. (A) Multi-dimensional scaling (MDS) plot to illustrate a 2D projection of differential gene expression between samples. Samples collected at different time points during infection (PI 7-27) as well as the parasites cultured with the medium are indicated by different colors. Two replicates are marked as A and B, respectively. (B) Sample hierarchical analysis heat map using Pearson correlation. The levels of difference between samples are indicated by different colors. (C) Venn diagram to indicate *L. passim* genes up-regulated at four different time points (PI 7-27) during infection (*P* < 0.01). (D) Venn diagram to indicate *L. passim* genes down-regulated at four different time points (PI 7-27) during infection (*P* < 0.01).

Listed genes specifically up-regulated at PI 7 (262) were enriched with GO terms such as “proton transmembrane transport” and encode several proteins associated with the kinetoplast electron transport chain, such as ATPase subunit 9 and cytochrome c oxidase subunits VI and X. GO-terms “cytoplasmic translation”, “nucleosome assembly” and “catalase activity” were enriched within the genes specifically down-regulated at PI 7. Many ribosomal protein-coding genes and genes associated with nucleosome assembly were down-regulated (Supplementary Table 4). These results suggest exposure of *L. passim* to the anaerobic and poor nutritional environment of the honey bee hindgut (compared to culture medium) stimulates oxidative phosphorylation and decreases protein translational activity by suppressing ribosome biogenesis. Many genes were up- (591) or down-regulated (575) specifically at PI 12 (Supplementary Table 5), with enrichment of only a few GO terms for the down-regulated genes, namely “cellular carbohydrate metabolic process” and “nucleoside salvage”. The genes continuously up-regulated at PI 12-27 (168) and PI 20-27 (126) encode proteins with various functions without specific GO enrichment (Supplementary Table 6). However, 463 genes continuously down-regulated at PI 12-27 and PI 20-27 included amastin surface glycoproteins, paraflagellar rod proteins, and inner kinetoplast membrane proteins such as cytochrome oxidase subunit IX (Supplementary Table 7). Indeed, we found that two GO terms, “inner mitochondrial membrane protein complex” and “integral component of membrane,” were enriched. Identification of differentially expressed genes (DEGs) between the sequential stages of *L. passim* infection demonstrated that the amount of various ribosomal protein mRNAs increased from PI 7 to PI 12 followed by a decrease at PI 20. Interestingly, many genes associated with rRNA synthesis and processing were down-regulated from PI 7 to PI 12 (Supplementary Table 8).

### Changes in the gene expression profile of the honey bee hindgut upon *L. passim* infection

We next compared the gene expression profiles between *L. passim*-infected and uninfected control honey bee hindguts (rectums) at the same ages (PI 7-27) by RNA-seq. The mapping rates of clean reads derived from control honey bees to the honey bee genome were 75-92 % and those derived from *L. passim*-infected honey bees were 8-68 %. The mapping rates were highest at PI 7, the time point with the lowest level of *L. passim* infection. Clustering analysis revealed that the overall gene expression profiles of the control honey bee hindguts were similar to those of the *L. passim*-infected ones, except for PI 12 (Fig. 3A and B) as for *L. passim* gene expression profile (See above). These results indicate that gene expression in the honey bee hindgut does not change dramatically upon *L. passim* infection. In fact, the number of DEGs between *L. passim*-infected and the control honey bee hindguts was in a range of 21 (PI 7) to 685 (PI 12) and only a few of them were shared between different infection time points (Fig. 3C and D) (Supplementary Table 9). Honey bees increased the expression of two antimicrobial peptide (AMP) mRNAs, *hymenoptaecin* and *defensin-1*, at PI 7. However, *inhibitor of dorsal* (*Cactus 1*) mRNA increased and another AMP, *abaecin*, mRNA was down-regulated. Thus, *L. passim* infection induces some but not all honey bee innate immune pathways. Interestingly, mitochondrial DNA encoding genes for cytochrome c oxidase subunit, ATP synthase F0 subunit, NADH dehydrogenase subunit, and cytochrome b, were up-regulated at PI 12. 589 genes associated with RNA synthesis and processing were down-regulated (Supplementary Table 10). The increased number of *L. passim* in the honey bee hindgut at PI 12 appears to enhance oxidative phosphorylation and suppress RNA synthesis and processing in the gut epithelial cells. Among the few genes regulated at the multiple stages of *L. passim* infection, immune system-related genes, *defensin-1* and *β-1,3-glucan-binding protein 1* (*βGBP1*) were up-regulated in PI 7-20 and PI 12-20, respectively. Similarly, nutrition and starvation-related genes, *take-out-like carrier protein* (*TO*) and *facilitated trehalose transporter Tret1-like* (*Tret1*) were up-regulated in PI 12-27 and PI 7-20, respectively. These results suggest that *L. passim*-infected honey bees are likely to be starved and that the gut epithelial cells are in poor nutritional status.

**Figure 3.**
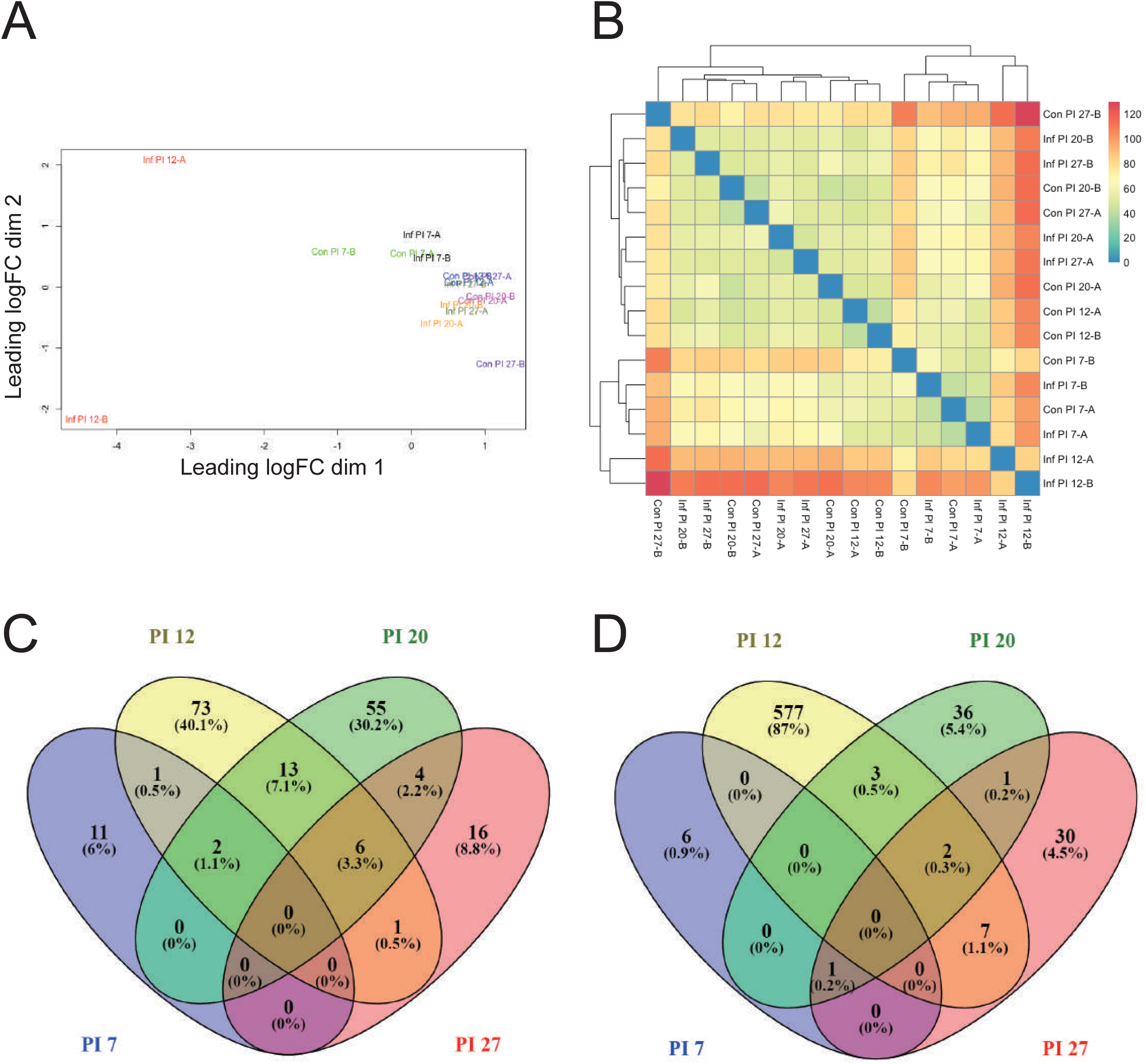
Global transcriptomic profiles of honey bee hindguts at different time points during infection by *L. passim*. (A) MDS plot to illustrate a 2D projection of gene expression differences between samples. *L. passim*-infected (Inf PI 7-27) and the uninfected control (Con PI 7-27) samples at different time points during infection are indicated by different colors. Two replicates are marked as A and B. (B) Hierarchical analysis heat map using Pearson correlation. The levels of difference between samples are indicated by different colors. (C) Venn diagram to indicate honey bee genes up-regulated in response to *L. passim* infection at four different time points (PI 7-27) (*P* < 0.05). (D) Venn diagram to indicate honey bee genes down-regulated in response to *L. passim* infection at four different time points (PI 7-27) (*P* < 0.05).

### Change of *Vitellogenin* (*Vg*) mRNA in the fat bodies of *L. passim*-infected honey bees

Enhanced expression of *TO* and *Tret1* mRNAs in the hindguts of *L. passim*-infected honey bees led us to test *Vg* mRNA levels in their fat bodies. *Vg* mRNA levels were shown to be correlated with the nutritional status of worker honey bees (Di Pasquale et al. 2013). As shown in Fig. 4, *Vg* mRNA levels in the fat bodies of honey bees at 20 days after *L. passim* infection were significantly lower than those of uninfected controls. These results suggest that *L. passim* infected-honey bees are in poor nutritional status.

**Figure 4.**
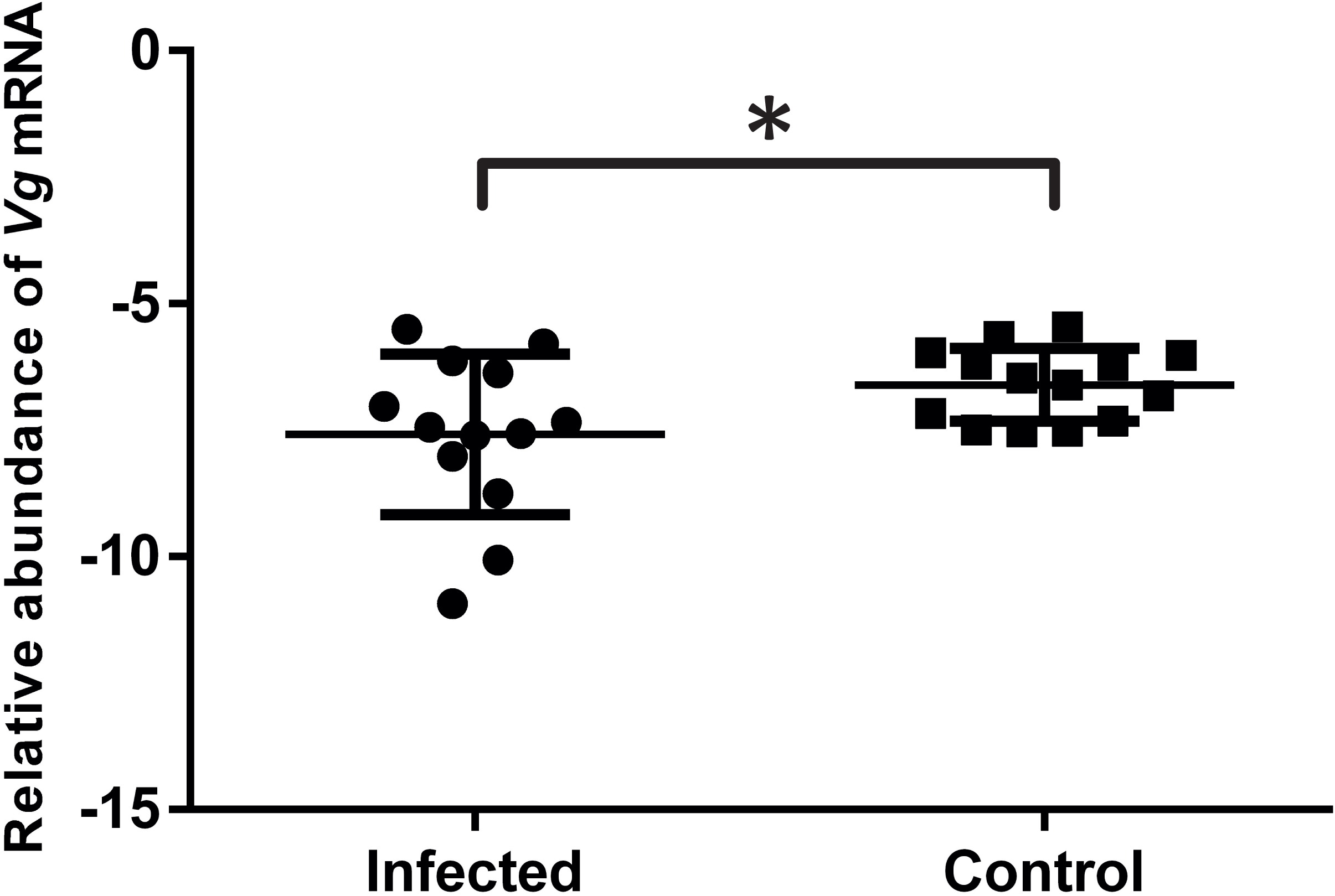
Relative *Vg* mRNA expression in the fat bodies of *L. passim*-infected and uninfected control honey bees. Relative expression of *Vg* mRNAs in the fat bodies of 13 *L. passim*-infected and 14 uninfected control honey bees was measured at 20 days after infection under natural conditions. Samples were obtained from two independent experiments. Mean values ± SD (error bars) are shown. Unpaired *t*-test (two-tailed) was used for statistical analysis (* *P* < 0.05).

## Discussion

### Effects of *L. passim* infection on the honey bee

We confirmed that ingested *L. passim* reaches the honey bee hindgut and colonizes it as previously reported (Schwarz et al. 2015). The number of parasites dramatically increases around 7-12 days and then reaches a plateau at 20 days after infection with 10^5^ cells under both laboratory and hive conditions. Accumulation of the parasites in the honey bee hindgut slightly increases bee mortality and never results in rapid death. This appears to be the case under hive conditions as well, since we recovered *L. passim*-infected honey bees at 37 days after infection. Many parasites share these infection characteristics in order to increase dissemination to other individuals through feces (Higes et al. 2016, Koch and Schmid-Hempel 2011, Koch et al. 2017, Langridge and McGhee 1967).

We did not find evidence for interactions between the honey bee gut microbiota (Firmicutes and universal bacteria) and *L. passim.* The number of gut bacteria was not affected by the *L. passim* infection and a similar observation had been previously made (Hubert et al. 2017). Interestingly, *L. passim* infection increased the number of *Gilliamella apicola* and universal bacteria in hives but not under laboratory conditions with a higher level of infection (Schwarz et al. 2016). This may suggest that a small number of *L. passim* modifies the honey bee hindgut environment and stimulates colonization of gut microbiota but a high degree of modification by a large number of parasites does not support further colonization. Lack of interaction between *L. passim* and honey gut microbiota is also consistent with recent studies that show the bumble bee microbiota is unaffected by *C. bombi* exposure and infection (Mockler et al. 2018, Näpflin and Schmid-Hempel 2018). Nevertheless, whether *L. passim* affects the number of other bacterial species that were not examined in this study and/or the functions of the gut microbiota - rather than just the number - remains to be tested.

### Dynamic changes in the gene expression profile of *L. passim* during honey bee hindgut infection

We found that *L. passim* dramatically changes its gene expression profile throughout the different stages of infection in the honey bee hindgut. Many ribosomal protein-coding genes are first down-regulated at the early stage of infection (PI 7), up-regulated at the middle stage (PI 12), and then down-regulated again at the late stage (PI 20-27). The nutritional status of the honey bee hindgut must be poor compared to that of culture medium because most of the nutrients derived from pollen and honey (food) are absorbed in the midgut and only residual compounds of food and metabolites from the gut microbiota are available. Thus, limited supply of nutrients may down-regulate the ribosomal protein-coding genes to suppress protein translation at first. However, *L. passim* adapts to the nutritional status of the honey bee hindgut, starts proliferating, which can be inferred by the increase in the expression of ribosome-coding genes, and then stops dividing as it decreases expression of ribosome-coding genes. Lack of oxygen in the honey bee hindgut appears to have a dramatic effect on *L. passim* gene expression and metabolism as well. *L. passim* was initially cultured under normoxia and migrated into the anaerobic honey bee hindgut, causing it to up-regulate the genes involved in the kinetoplast electron transport chain to compensate for the lack of oxygen at the early infection stage (PI 7). However, these genes appear to be down-regulated at later infection stages as the parasite switches to a dormant stage without active metabolism and proliferation. Similar shifts in energy metabolism are also observed in other trypanosomatid parasites (Bringaud et al. 2006, Smith et al. 2017, Van Grinsven et al. 2009, van Hellemond et al. 2005). Furthermore, the continuous down-regulation of *tryparedoxin 1, tryparedoxin-like*, and *tryparedoxin peroxidase* genes throughout the infection cycle is also consistent with the reduced production of radical oxygen species in anaerobic environments. Thus, *L. passim* appears to shift its growth and metabolism depending on the nutritional and anaerobic conditions of the honey bee hindgut.

In terms of morphology, *L. passim* grown in culture medium presents a flagellum. However, the flagellum is absent when the parasites accumulate in the honey bee hindgut. This is also supported by the down-regulation of genes associated with flagellar formation such as paraflagellar rod proteins (Portman and Gull 2010). *L. passim* appears to change its cell surface proteins during the infection cycle. *Amastin* surface glycoprotein genes are down-regulated during middle to late stages of infection and *GP63* is instead up-regulated throughout the infection cycle. The functions of amastin were tested in *Leishmania* and it was described as necessary for the efficient interaction between amastigotes and the membrane of parasitophorous vacuoles in the infected macrophage (de Paiva et al. 2015). *Leishmania* GP63 was proposed to be essential for the promastigotes to attach to the insect gut wall (D’Avila-Levy et al. 2006, d’Avila-Levy et al. 2014, de Assis et al. 2012). *Leptomonas* GP63 was suggested to have the same function (Pereira et al. 2009), which may also apply to *L. passim* GP63. Furthermore, *Leishmania* GP63 was shown to activate host protein tyrosine phosphatases that lead to the repression of several protein kinase signaling pathways (Blanchette et al. 1999, Gomez et al. 2009, Shio et al. 2012). Thus, up-regulation of *L. passim* GP63 could be critical to establish and maintain the infection in the honey bee hindgut.

### Responses of the honey bee host to *L. passim* infection

The honey bee hindgut response to *L. passim* infection is characterized by changes in gene expression. However, the number of DEGs is relatively small and very few of them are shared at different infection time points. Similar to the *C. bombi* infection in bumble bee, we found that two *AMPs, defensin-1*, and *hymenoptaecin*, are induced by the *L. passim* infection in the honey bee hindgut. Since these bumble bee AMPs were shown to directly inhibit the growth of *C. bombi* (Deshwal and Mallon 2014, Marxer et al. 2016), we could also expect they suppress the growth of *L. passim*. However, we also found that *abaecin* was down-regulated at PI 7 and this could be the result of a Toll signaling pathway suppression caused by the increase of *Cactus* (*inhibitor of dorsal*) mRNA. Based on the metabolic pathways identified by genome sequencing, *L. passim* should be able to synthesize β-1,3-glucan on its cell surface, like fungus, and βGBP1, which was upregulated in the honey bee hindgut, could bind to *L. passim* through glucan moiety (Söderhäll and Unestam 1979). This would result in the activation of prophenoloxidase (pro-PO) (for melanin deposit) or the Toll signaling pathway (Brutscher et al. 2015, Cerenius et al. 2008, Honti et al. 2014, Tang et al. 2006). *Defensin-1* and *βGBP1* were up-regulated at multiple stages of the *L. passim* infection (PI 7-20 and PI 12-20, respectively), suggesting that the honey bee appears to fight the infection by activating its innate immune system. It is surprising that transcription of multiple mitochondrial DNA-encoded genes was stimulated only at PI 12 without simultaneous up-regulation of nuclear DNA-encoded genes associated with the electron transport chain. As a result, the overall activity of oxidative phosphorylation may not change very much at PI 12. Increase of *L. passim* in the hindgut lumen may transiently inhibit oxygen supply to gut epithelial cells from the basolateral side by, for example, breaking tight junctions. TO was shown to be involved in controlling the feeding behavior and nutritional status of fruit fly by binding juvenile hormone and mRNA expression is highly stimulated in the head and gut tissues by starvation (Sarov-Blat et al. 2000, So et al. 2000). Thus, up-regulation of *TO* in the hindgut at PI 12-27 suggests that *L. passim*-infected honey bees were under starvation. Trehalose levels in the hemolymph are reduced in the fruit fly under starvation (Yamada et al. 2018) and this is consistent with the increase of *Tret1* mRNA in the hindgut of *L. passim*-infected honey bees at PI 7-20. The decrease in hemolymph trehalose levels is caused by its reduced release from fat bodies and increased uptake by various tissues. Accordingly, we also found *Vg* mRNA is reduced in the fat bodies of *L. passim*-infected honey bees compared to the uninfected controls. Honey bee gut microbiota was shown to produce various metabolites such as organic acids, some of which would be absorbed by the host and affect weight gain as well as appetite behavior (Zheng et al. 2017). *L. passim* may suppress this process and lead the honey bee into a poor nutritional status.

Our results show how *L. passim* modifies its gene expression to adapt to the honey bee hindgut environment and, at the same time, how the honey bee modifies its gene expression against the *L. passim* infection. This study describes one of the best models to understand host (honey bee)-parasite (*L. passim*) interactions at the molecular and cellular levels.

## Materials and Methods

### *L. passim* and honey bee sample preparation

To prepare for the infection in honey bees, *L. passim* cells were grown in modified FPFB medium (Salathe et al. 2012) supplemented with 10 kU/mL penicillin (Beyotime), 10 mg/mL streptomycin (Beyotime), and 50 mg/mL gentamycin (Noble Ryder) at 25 °C. The cells were collected during logarithmic growth phase and washed once with PBS followed by resuspension in sterile 10 % sucrose/PBS at 20, 000 viable cells/μL. Newly emerged honey bee workers were collected and fasted for 2-3 h. Sucrose solution was given to confirm most of them were starved for inoculum intake. Honey bees were divided into two groups: One group was fed with 5 μL of 50 % sucrose solution (Control). The other group was fed with 5 μL of 50 % sucrose solution containing *L. passim* (100,000 cells in total). After the oral infection, the honey bees were marked with oil paint on the thorax and returned to the hive. Alternatively, they were kept in metal cages at 33 °C under laboratory conditions.

### Quantitative PCR (qPCR) to analyze the abundance of *L. passim* in the infected honey bee guts

Ten honey bees were collected at 1, 3, 8, 15 and 22 days after infection and genomic DNA was extracted from the whole abdomens of individual bees using DNAzol® reagent (Thermo Fisher). To quantify *L. passim* infection levels, the *internal transcript spacer region 2 (ITS2)* of *ribosomal RNA (rRNA)* was used as a target for PCR amplification. Honey bee *AmHsTRPA* was used as the internal reference. The relative abundances of *L. passim* in the individual honey bees were calculated by the ΔC_t_ method. The Steel-Dwass method was used for statistical analysis.

### Mortality test for *L. passim*-infected honey bee workers

To test the effects of *L. passim* infection on the mortality of honey bees, we kept 100 *L. passim*-infected and control worker honey bees in metal cages and fed them with 50 % sucrose solution at 33°C. The dead bees were counted and removed from the cages every day. The mortality test was repeated three times. The results were statistically analyzed by the Log-rank (Mantel-Cox) test. In order to confirm *L. passim* infection, dead bees (one from the control and six from the *L. passim*-infected group) were sampled 15 and 16 days after infection for each experiment. Genomic DNA was extracted from individual bees and the parasite was detected by PCR of the *rRNA ITS1* (Da Silva et al. 2004). Honey bee *AmHsTRPA* was used as a control. Six dead bees from the infected groups were specifically positive for *L. passim*.

### Quantification of the relative abundance of universal bacteria and Firmicutes in honey bee guts under laboratory conditions

Newly emerged honey bee workers were kept in frames for 48 h for acquisition of the core gut microbiota. The honey bees were then divided into an *L. passim*-infected group and a control group, as above. The honey bees were fed with 50 % sucrose solution together with a pollen mixture. We collected eight bees from each group at 7 and 15 days and seven bees at 22 and 27 days after infection. Bee abdomens were washed with 75 % ethanol followed by immersion in sterilized PBS. Whole guts were isolated and genomic DNA was extracted. Primers described in previous reports (De Gregoris et al. 2011, Powell et al. 2014) were used to quantify universal bacteria and phylum-specific *Firmicutes*. The honey bee *β*-*actin* gene was used as a reference (Li et al. 2017). The statistical analysis was carried out by unpaired *t*-test (two-tailed).

### Preparation of *L. passim* and honey bee samples for RNA-seq

*L. passim*-infected and control honey bees were sampled from the hive at 7, 12, 20, and 27 days after infection. Total RNA was extracted from the rectums of more than two honey bees for each time point. The degree of *L. passim* infection was determined by quantifying *glyceraldehyde 3-phosphate dehydrogenase* (*GAPDH*) mRNA (Kojima et al. 2011) by qRT-PCR. Honey bee *elongation factor 1-alpha* (*Ef-1alpha*) mRNA was used as reference to verify for the quality of the RNA extraction as well as that of the reverse transcription (Kojima et al. 2011). Two samples with similar levels of *L. passim* infection were selected at each time point for RNA-seq. Total RNA was also extracted from *L. passim* cultured at the logarithmic growth phase. All of the above samples were sequenced using an Illumina HiSeq-4000 platform at the Beijing Genomics Institute (BGI). At least 4GB of clean data were obtained from each sample. The RNA-seq data were deposit to SRA database with the accession number of PRJNA510495.

### Analysis of the RNA-seq data of honey bee and *L. passim*

Honey bee reference genome assembly (GCF_000002195.4_Amel_4.5_genomic.fna), annotation GFF (GCF_000002195.4_Amel_4.5_genomic.gff) and protein fasta (GCF_000002195.4_Amel_4.5_protein.faa) files were downloaded from NCBI. The program gffread (http://ccb.jhu.edu/software/stringtie/gff.shtml) was used to convert the honey bee GFF file to GTF for HISAT2-build on a local MacBook Pro. The splice junctions and the positions of exons were extracted from the converted GTF file by using hisat2_extract_splice_sites.py and hisat2_extract_exons.py scripts from HISAT2 (Version: 2.1.0) (Kim et al. 2015). Next, HISAT2-build was used to construct the HISAT2 index with the spliced sites and exons. Clean reads were then mapped to the latest version of the honey bee index, and SAM files were converted and sorted by SAMtools (Li et al. 2009). HTSeq-count was used to count from each sorted SAM file. The NCBI RefSeq name was used as a unique identifier to count the reads from SAM files. The genome annotation of the honey bee was downloaded in tabular format to align the RefSeq name identifier to the functional annotation. Similarly, the genomic sequence of *L. passim*, strain SF (GenBank number: AHIJ00000000.1) was downloaded from NCBI (Runckel et al. 2014) to build the HISAT2 index. The annotation GFF file used for HTSeq-count was generated from Companion pipeline (Steinbiss et al. 2016) as described below. The protein fasta file from Companion pipeline was used to annotate the function of each protein-coding gene.

### Reannotation of the *L. passim* genome sequence by Companion pipeline

The clean reads of RNA-seq were mapped to *L. passim* genome using TopHat V2.1.0 with default parameters (Trapnell et al. 2009). All transcripts containing the *L. passim* genome sequence were assembled using Cufflinks v2.2.1 (Trapnell et al. 2012). Once the short read sequences were assembled, all transcripts were merged using Cuffmerge (Trapnell et al. 2012). The merged *L. passim* transcript and genomic sequences were annotated using the parasite genome annotation pipeline Companion with *Leishmania major Friedlin* as the reference organism (Steinbiss et al. 2016). In the pseudochromosome contiguation, the minimum length required for contig placement was set at 200 bp and the minimum similarity for contig placement was set at 35 %. The reannotated GFF3 file and the protein coding sequences were used for the Differential Gene Expression (DGE) and other downstream analyses.

### DGE analysis of both *L. passim* and honey bee

Raw read counts from both *L. passim* and honey bee were analyzed in RStudio (Version 3.4.3) using the generalized linear model (GLM)-based method of edgeR package (Version 3.20.9) from Bioconductor (Robinson et al. 2010). Normalization was performed with Trimmed Mean of M-values (TMM) implemented in the edgeR Bioconductor package (Robinson and Oshlack 2010). A filter was set up to remove any reads with less than one count per million mapped reads (CPM). After testing, the Benjamini-Hochberg (BH) method was applied to control the false discovery rate (FDR) across the detected loci. For *L. passim,* the raw *P*-values for each gene were corrected and the differentially expressed genes (DEGs) were identified by the threshold of FDR < 0.01. For the honey bee, the DEGs were identified by the threshold of FDR < 0.05. A multi-dimensional scaling (MDS) plot and Pearson correlation-based heat maps created by the gplots pheatmap.2 function were used to show the distances between samples. After obtaining the DEGs, Venny (http://bioinfogp.cnb.csic.es/tools/venny/index.html) was used to build the Venn diagrams to summarize the relationships between DEGs identified in *L. passim* and in the honey bee under different conditions.

### Gene ontology (GO) enrichment analysis (Fisher’s Exact Test)

In order to reduce the time for computational analysis and increase the specificity of the study, local blast databases were made using Blast2Go software with the taxonomy IDs. In total, 446,413 protein sequences from the *Order Kinetoplastida* (Taxonomy ID: 5653) and 6,225,195 protein sequences from the *Subphylum Hexapoda* (Taxonomy ID: 6960) were downloaded from the NCBI database with GI numbers. All of the above sequences were concatenated into two FASTA files separately. These two FASTA files were then used to build the local BLAST database in the Blast2GO PRO desktop version 5.0.1 (Conesa et al. 2005). After obtaining the DEGs from *L. passim* and the honey bee, the up- and down-regulated genes were analyzed individually at different time points. For each data set, protein sequences were extracted from the whole protein FASTA file of *L. passim* or *A. mellifera* using python script and imported to Blast2GO. Local BLAST was then performed using blastp (parameters: E-Value Hit Filter = 1e^-10^; Number of Blast Hits: 20) against the appropriate local BLAST database created above. For each dataset, an InterPro scan was simultaneously run using the EMBL-EBI public web-service database with the default settings. The mapping and annotation procedures were carried out using the latest online databases according to default settings. After annotation, the InterProScan GOs were merged and the enrichment analysis (Fisher’s Exact Test) was performed using a program embedded in Blast2GO. The entire protein-coding sequences of *L. passim* and *A. mellifera* were used to generate the reference annotation lists for the GO enrichment analysis. After obtaining the enriched GO terms, a cut-off of FDR < 0.05 was applied to list the most specific GOs.

### Quantification of *Vitellogenin (Vg)* mRNA in the fat bodies of *L. passim*-infected and the control honey bees by qRT-PCR

*L. passim*-infected and control honey bees were collected from the hive at 20 days after infection. The fat bodies of individual bees were dissected, and then total RNA was extracted as described above. Expression of *Vg* mRNA relative to *18S rRNA* was measured by qRT-PCR (Ihle et al. 2015) and calculated using the ΔC_t_ method (Schwarz et al. 2016). The results were statistically analyzed by unpaired *t*-test (two-tailed). Sequences of all primers used for PCRs are listed in Supplementary Table 11.

